# Antagonistic roles of Polycomb repression and Notch signaling in the maintenance of somatic cell fate in *C. elegans.*

**DOI:** 10.1101/055137

**Authors:** Ringo Pueschel, Francesca Coraggio, Alisha Marti, Peter Meister

## Abstract

Reprogramming of somatic cells in intact nematodes allows characterization of cell plasticity determinants, which knowledge is crucial for regenerative cell therapies. By inducing muscle or endoderm transdifferentiation by the ectopic expression of selector transcription factors, we show that cell fate is remarkably robust in fully differentiated larvae. This stability depends on the presence of the Polycomb-associated histone H3K27 methylation, but not H3K9 methylation: in the absence of this epigenetic mark, many cells can be transdifferentiated which correlates with definitive developmental arrest. A candidate RNAi screen unexpectedly uncovered that knock-down of somatic Notch^LIN-12^ signaling rescues this larval arrest. Similarly in a wild-type context, genetically increasing Notch^LIN-12^ signaling renders a fraction of the animals sensitive to induced transdifferentiation. This reveals an antagonistic role of the Polycomb repressive complex 2 stabilizing cell fate and Notch signaling enhancing cell plasticity.

## Introduction

During development, cellular potency is progressively restricted and differentiated cells have lost their plasticity. *C. elegans* conforms to this paradigm: early embryonic blastomeres can be converted in a number of cell types by ectopically expressing selector transcription factors^1–5^, while later during development most cells lose this capacity. In fully differentiated animals, only one transcription factor, the endodermal-specifying ELT-7 was shown to be able to induce transdifferentiation of pharyngeal cells into an intestinal-like cell type^6^. Nematodes are an interesting system to characterize the molecular players modulating somatic cell fate plasticity during development^7^. Previous studies showed that in embryos, the elimination of the Polycomb complex or GLP-1^Notch^ signaling extend the blastomeres plasticity period^8,9^. In differentiated animals, only few factors are known to modulate cell plasticity, most of which were characterized in a natural transdifferentiation event, the endodermal Y to neuronal PDA conversion^10–12^. Chromatin modifications appear prominent, as the temporally controlled expression of distinct histone modifiers is necessary for conversion^11^. Here we report a singlecopy cell fate induction system for muscle and endoderm. Using muscle induction, we show that cell fate is remarkably stable in fully differentiated animals of the first larval stage as only one cell was found to transiently express muscle markers. In contrast, in the absence of the Polycomb complex muscle fate induction leads to aberrant expression of muscle markers in numerous cells as well as a robust larval arrest. Using this arrest as a reporter for cell plasticity regulators, we demonstrate a role of the Notch signaling pathway in enhancing cellular plasticity, thus antagonizing Polycomb cell fate stabilization.

## Results and Discussion

All systems used to date to induce transdifferentiation are based on multicopy arrays (reviewed in ^13^). Arrays are epigenetically silenced and highly enriched for silent histone marks (histone H3 lysine 9 and 27 (H3K9/27) methylation)^14,15^. This renders the analysis of the function of these marks on cell plasticity difficult. To overcome this issue, we integrated single copies of a heat-shock (HS) inducible construct driving either a muscle^4^ (*hlh-1/MyoD*) or an endoderm^3^ (*end-1*/GATA1) specifying transcription factor. A trans-spliced *mCherry* ORF is placed downstream of the transcription factor sequence to control for expression (Figure 1A). Muscle cells are identified by expression of *gfp::histone* H2B controlled by the heavy chain myosin promoter *myo-3*. As for arrays, ectopic *hlh-1* or *end-1* expression in early embryos (20-100 cells) leads to irreversible developmental arrest (Figure 1B). Moreover, ectopic HLH-1 induced cellular twitching about 10 hours post induction and many (but not all) cells in these embryos displayed green nuclei, suggesting transdifferentiation into muscle fate (Figure 1B, HLH-1^ect^, arrows). As previously reported^8^, cellular plasticity is lost later during development, where expression of either transcription factors has no phenotypic effect and animals hatch normally (Figure 1-S1).

**Figure 1.**
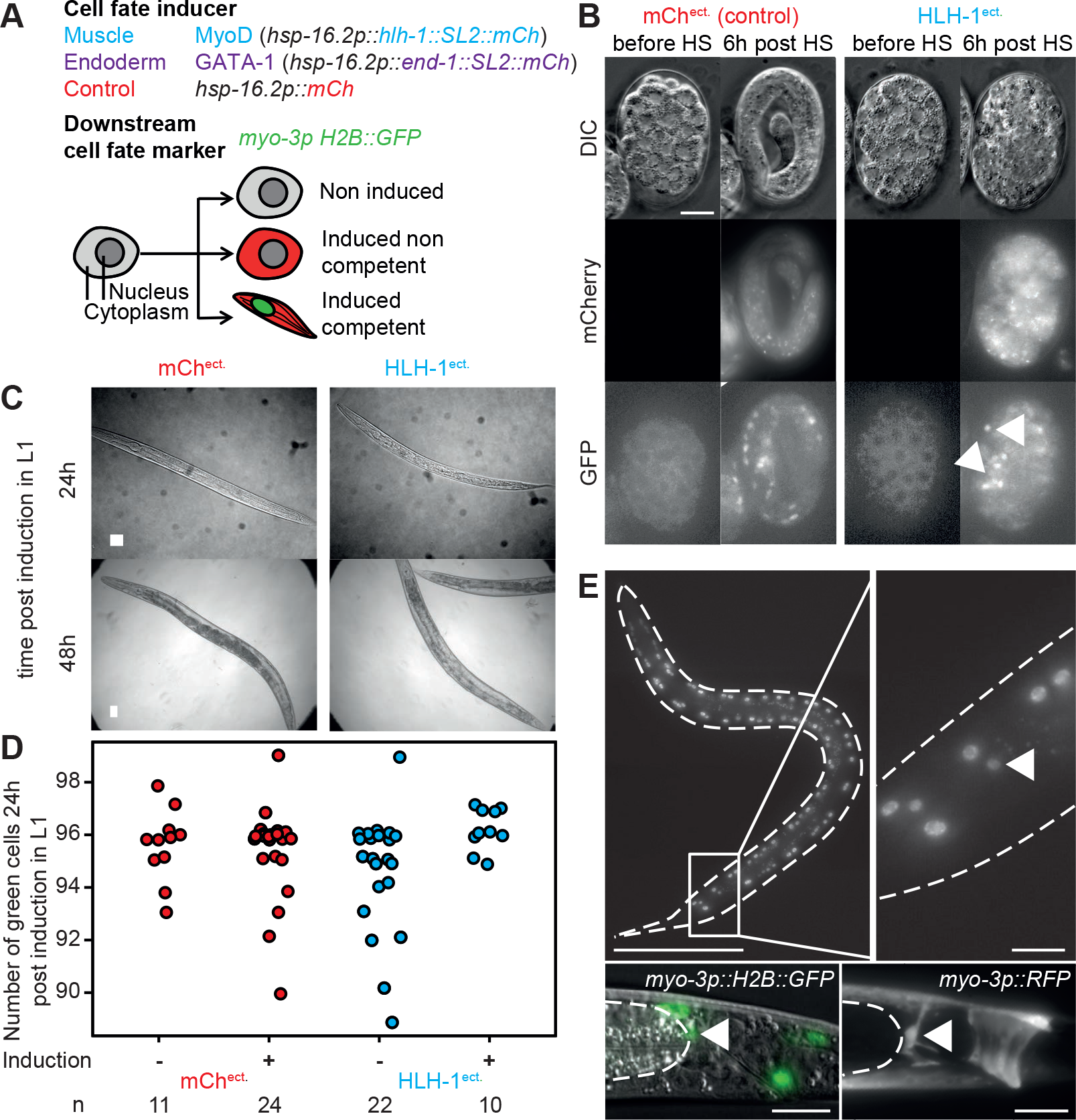
**A.** Single-cell readout cellular plasticity sensor. Cell-fate specifying transcription factors *hlh-1* (MyoD homolog, inducing muscle fate) or *end-1* (GATA-1 family homolog, inducing endodermal/intestinal fate) are induced by heat-shock treatment. Both transcription factors open reading frames are placed upstream of a trans-spliced SL2 *mCherry* ORF, providing a visual readout for their expression. A cell fate marker (histone H2B::GFP) for muscle fate is integrated elsewhere in the genome. All constructs are present as single copy insertions. Upon induction of the expression of the transcription factor, red cytoplasmic fluorescence indicates correct induction while nuclear green fluorescence informs on the muscle differentiation potential of the given cell. **B.** Muscle cell fate induction in early embryos (~35 cell stage). Pictures show DIC, red and green fluorescence channels before induction and 6h post induction, in a *mCherry* control strain and upon HLH-1 ectopic expression. Upon expression of HLH-1, embryos rapidly arrest development and a number of cell express the muscle-specific marker (arrows). Scale 10 μm. **C.** Brightfield images of wild-type worms ectopically expressing either *mCherry* or *hlh-1* 24 and 48 hours post induction. Bar 25 μm. **D.** Number of GFP::H2B positive cells of non-induced and induced worms 24 hours post induction. Red: *mCherry* control; blue: HLH-1. **E.** Upper left: GFP::H2B signal in an animal ectopically expressing HLH-1 imaged 24 hours post induction (z maximal intensity projection). Bar 100 μm. Upper right: magnification of the tail region. The additional nucleus expressing GFP::H2B is indicated with an arrow. Bar 10 μm. Lower left: DIC/green fluorescence overlay of the tail of an animal of the same strain, imaged 24 hours post induction of HLH-1. Bar 10 jm. Dashed line: gut, white arrow: additional cell expressing muscle markers. Lower right: same tail region than in lower left, imaged 24 hours post HLH-1 induction in a strain carrying a cytoplasmic red muscle marker *(myo-3p::RFP)*. Bar 10 μm. The cytoplasmic RFP signal outlines the characteristic shape of the anal-sphincter cell (arrow).

## Cell fate of differentiated animals is robust

The fluorescent muscle fate reporter allows the visualization of potentially transdifferentiating cells beyond embryonic development. When HLH-1 was expressed in fully differentiated first larval stage animals, twenty-four hours post induction (animals in L2-L3 stage), about 50% of the animals had one more than the normal 96 *myo-3* expressing cells (Figure 1D). This additional cell was located in the tail region between the gut and the rectum and present in 46% of the animals (n=122), but never observed in control HS larvae (n=91). This location corresponds to the anal sphincter cell^16^ (Figure 1E, arrow). Indeed, a cytoplasmic *myo-3p::RFP* marker highlighted the typical, saddle-like shape of this cell in 9 out of 13 worms 24h post *hlh-1* induction (Figure 1E, bottom right). 48 hours post induction however, high expression of the muscle marker in the anal sphincter cells was no longer visible (Figure 1-S2). We conclude that upon HLH-1 expression, the anal sphincter cell transiently expresses muscle-specific markers but subsequently represses these, supposedly reverting to its normal fate. Interestingly, this cell is epigenetically close to a muscle as its sister is a body wall muscle cell^16,17^. Reversion to silencing of the muscle marker might be a consequence of signaling from surrounding cells inhibiting complete fate conversion^18^. Altogether, our experiments demonstrate that differentiated animals are remarkably robust to induction of muscle transdifferentiation, with a single cell transiently expressing a muscle-specific marker.

## Absence of H3K9 methylation or perinuclear H3K9me anchoring has no effect on cell plasticity in differentiated animals

Anchoring of H3K9 methylated heterochromatin at the nuclear periphery was recently shown to help stabilize ectopically induced cell fates in embryos^19^. We therefore asked whether this feature could be extended to the first larval stage by testing mutants deficient for either H3K9 methylation or anchoring of methylated H3K9 at the nuclear periphery (*set-25 met-2* or *cec-4* mutants, respectively)^14,19^. In both mutants, ectopic expression of *hlh-1* did not lead to obvious phenotypical alterations nor did it increase the total number of green nuclei per worm or the proportion of animals in which the anal sphincter cell expressed muscle markers (data not shown and Figure 1-S3). As for wild-type animals, muscle marker expression was no longer observed in this cell 48 hours post induction. In conclusion, ablation of H3K9 methylation or its perinuclear anchoring does not impact on cellular plasticity in fully differentiated animals.

## Absence of the Polycomb Repressive Complex leads to larval arrest upon *hlh-1*^MyoD^ or *end-1*^GATA-1^ expression

H3K27 methylation, deposited by the Polycomb Repressive Complex 2 plays a crucial role in the modification of the epigenetic landscape during development of worms and other organisms^20–22^. Ablation of PRC2 components leads to an elongation of the embryonic plasticity window and renders germline cells amenable to fate conversions^8,23^. While *mes-2(RNAi)* animals are sterile, second generation *mes-2* homozygous mutant animals develop normally, although sterile. This generation has no detectable H3K27 methylation, hence the mark is dispensable for cell fate specification under normal conditions^24^. However, ectopic expression of HLH-1 in first larval stage *mes-2* F2 animals has dramatic effects: 93% of the worms arrest larval development (Figure 2A, compare with 1C; scoring in 3A). Germline cell number confirmed that most animals arrest at the L1 stage (Figure 2B, *mes-2* HLH-1^ect.^, insert). Developmental arrest is a consequence of HLH-1 expression and not the heat-shock used for *hlh-1* induction as heat-treated *mes-2* control animals developed normally (Figure 2A, Figure 3A). Although motile and alive, the developmentally arrested animals remain small and die without resuming development seven to ten days post-induction. Physiologically, these animals show reduced pharyngeal pumping (75 pumps per minute (ppm) *versus* 185 ppm for wild-type or control *mes-2* animals). Similarly to HLH-1, ectopic expression of END-1 leads to developmental arrest in 60% of *mes-2* animals, although at later stages of development (Figure 2C, Figure 3A). Arrest is as stringent as for HLH-1^ect.^ since these animals die within 3-7 days without resuming development.

**Figure 2.**
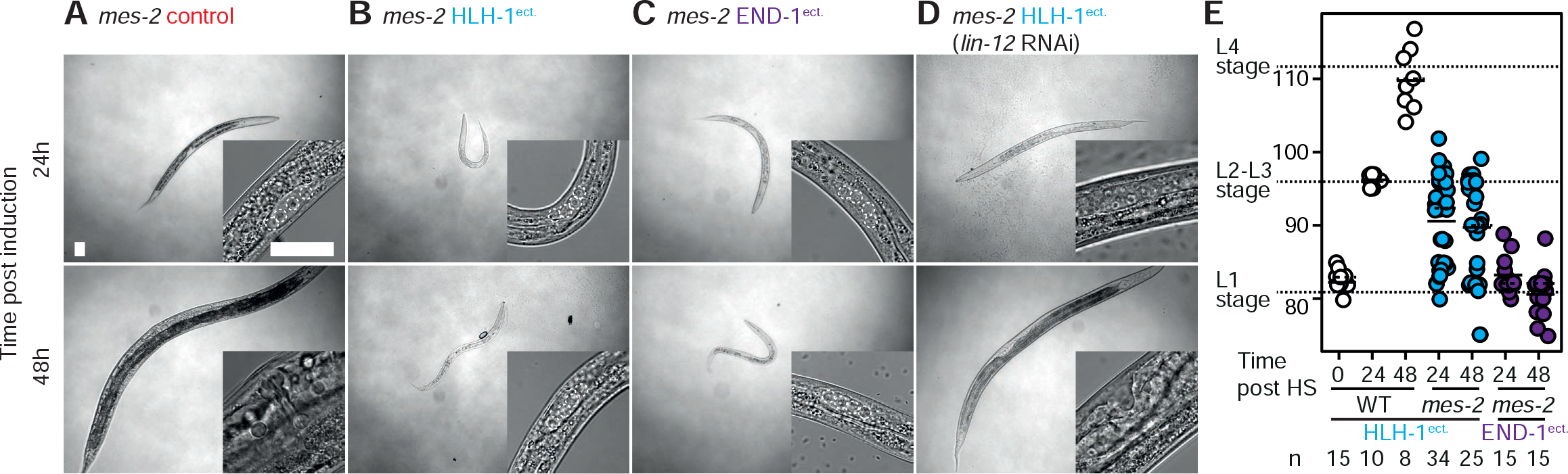
**A.** Brightfield images of *mes-2* mutant control animals 24/48 hours post heat-shock. Next to the picture of the entire animal (bar 25 μm), a magnification of the gonad/vulva is shown for staging purposes (bar 25 μm). **B.** Brightfield images of *mes-2* mutant animals ectopically expressing HLH-1 24/48 hours post induction. **C.** Brightfield images of *mes-2* mutant animals ectopically expressing END-1 24/48 hours post induction. **D.** Brightfield images of rescued *mes-2* mutant animals grown on *lin-12(RNAi)* ectopically expressing HLH-1 24/48 hours post induction. **E.** GFP::H2B positive nuclei in wild-type animals (white) and *mes-2* arrested animals upon ectopic expression of either HLH-1 (blue) or END-1 (purple) 24 and 48 hours post induction in the first larval stage of fed animals. The short solid line indicates the mean while the dashed line the median value.

**Figure 3.**
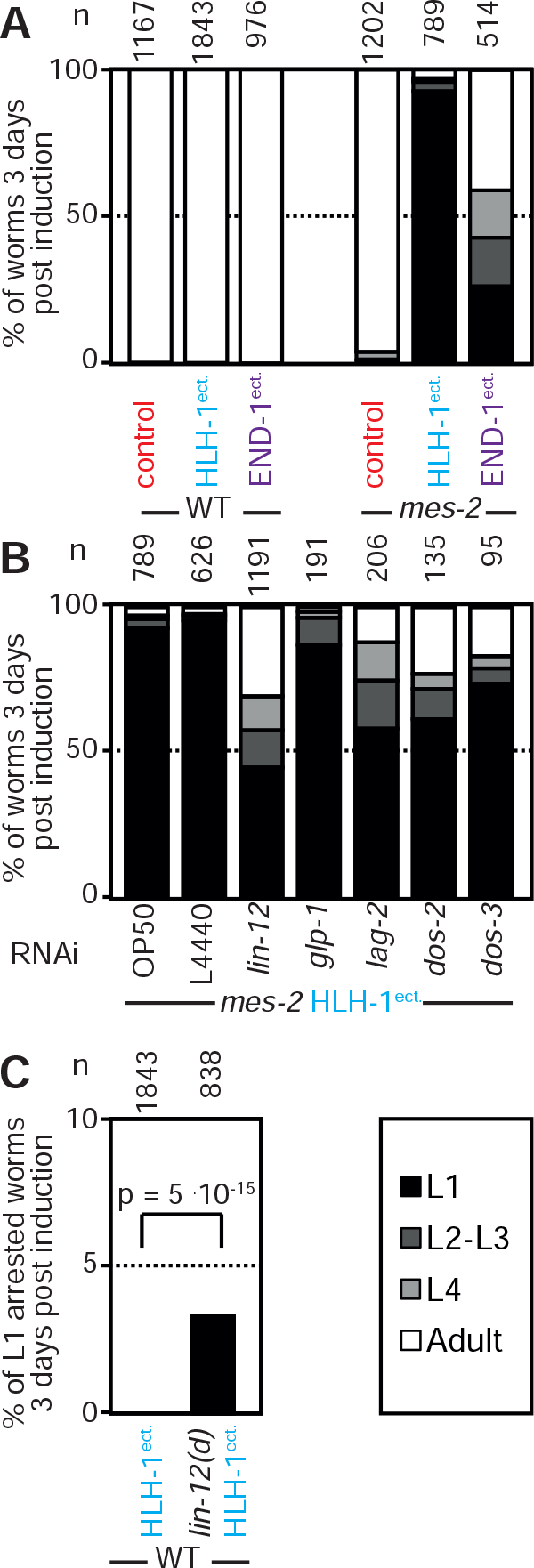
**A.** Scoring of animal development 3 days post induction at the first larval stage. Comparison of control animals to strains ectopically expressing HLH-1 or END-1 in a wild-type or *mes-2* genetic background. Proportions of the populations are shown. Black: animals in the first larval stage; dark grey: animals in the second/third larval stage; light gray: fourth larval stage; white: adult worms. **B.** Scoring of development 3 days post induction of HLH-1 expression at the first larval stage in *mes-2* animals fed on RNAi for the indicated genes. Color code as in A. **C.** Scoring of arrested L1 animals 3 days post induction of HLH-1 expression at the first larval stage in wild-type animals with the dominant *lin-12(n905)* mutation. Fisher Test p value 5.10^−15^.

In wild-type animals, apart from the anal sphincter cell, HLH-1^ect.^ expression has no effect on the number of cells expressing the muscle marker. This number increases from one larval stage to the next correlated with the number of muscle cells^16^ (Figure 2D, WT HLH-1^ect^). In contrast, more than 90% of the *mes-2* L1 arrested animals expressing HLH-1 have more than the 81 green nuclei per L1 animal, with numbers ranging from 82 to 97 (24h post HS: 33 out of 34; 48h post HS: 23 out of 25; mean cell number: 90.8, median: 90; Figure 2D *mes-2* HLH-1^ect^). In contrast, END-1^ect^. did not lead to the appearance of additional cells positive for the muscle marker but induced the same larval arrest suggesting it similarly perturbs cell fate (Figure 2D). Cells expressing the muscle marker upon HLH-1^ect^. did not show a preferential localization, unlike in wild-type animals. Moreover, RNA seq of arrested L1 animals failed to uncover a cell type specific transcriptional signature (data not shown). Collectively, this suggests that in the absence of PRC2, HLH-1^ect.^ and END-1^ect.^ induce stochastic transdifferentiation as observed in other systems^20,25,26^ and that this transdifferentiation leads to developmental arrest. Hence, although dispensable for normal fate specification, PRC2/H3K27me are essential to stabilize cell fate upon perturbations.

## Notch signaling enhances cell fate plasticity

If PRC2 is essential to terminate plasticity, knock-down of any factor enhancing plasticity could suppress larval arrest induced by the expression of HLH-1 or END-1 in *mes-2* animals. We therefore screened by RNAi a library of previously characterized genes involved in cell plasticity in *C. elegans* (Table S1). Most RNAi had no effect and a large majority of *mes-2* F2 animals arrested development upon HLH-1^ect^. expression (Figure 3-S1). In contrast, *lin-12(RNAi)* rescued 55% of the population which developed to adulthood *(versus* 96% animals arrested as L1 for control RNAi; Figure 3C). LIN-12 is one of the two Notch receptors homologs. Knock-down of the second homolog *glp-1* had no effect, in agreement with its major role in the germline and during embryogenesis^27^. Similarly to *lin-12* RNAi, knock-down of the Notch ligand components *lag-2, dos-2* and *dos-3* rescued larval arrest at a slightly lower degree than *lin-12* RNAi (42, 39 and 21% of animals reaching adulthood, respectively). Rescue from larval arrest by *lin-12(RNAi)* was also observed upon END-1^ect^. induction, in which we could detect a reduction from 18% to 10% of arrested L1 worms by *lin-12* RNAi (n=582, 664). We conclude that LIN-12^Notch^ signaling enhances cell plasticity, thereby antagonizing PRC2 stabilization of cell fate. Similarly, increasing LIN-12^Notch^ activity by introducing a gain-of-function *lin-12* allele in a PRC2 wild-type background leads to larval arrest upon HLH-1 expression in 3% (n=838) of the animals, while this was never observed in animals with normal Notch signaling (n=1843, p=5.10^−15^). Increasing Notch signaling therefore renders animals more sensitive to ectopic transcription factor expression. Interestingly, LIN-12^Notch^ signaling was previously shown to accelerate dauer exit, during which animals undergo major reorganization and cell fate changes^28^. Similarly, Notch signaling is essential to endow the endodermal Y cell with the competence for transdifferentiation and enhances germline cell reprogramming by antagonizing Polycomb-mediated silencing (see accompanying manuscripts). Together, this suggest a widespread mechanism in the germline and the soma to control cell fate plasticity by the fine balance between PRC2 restricting pluripotency and Notch signaling enhancing it.

## Methods

### General worm methods

Unless otherwise stated, *C. elegans* strains were grown on NG2 medium inoculated with OP50 bacterial strain at 22.5°C. The dominant *lin-12(n905)* gain of function mutation was introduced as in ^29^.

### Synchronization and TF induction in embryos and larvae

For embryos, wild-type or first generation *mes-2* gravid adults were dissected in M9, 1 and 2 cell stage embryos were transferred to the 2% agar pad and incubated at 22.5°C until they reached the desired stage as described in ^30^. The developmental stage was verified before HS by imaging and the expression of the transcription factor was induced by 10’ heat shock at 33°C in a PCR thermocycler.

For larvae synchronization, wild-type worms were synchronized either by letting gravid adults lay eggs for 2-4 hours on plates or by bleaching gravid adults. Embryos were then left to hatch overnight at 22.5°C. *mes-2* animals are sterile in the second generation. Synchronized F2 animals were prepared by manually picking F1 homozygotes (phenotypically Unc) from balanced parents (wild-type). F2 homozygotes were obtained as for the wild-type animals. For TF induction, synchronized worms were washed off the plates with M9, transferred to an Eppendorf tube, spun at 1000 g for 1 minute and washed once before spinning them again. The supernatant was then aspirated to concentrate the worms in a small volume (~20-50μL). Animals were heat-shocked in a 33°C water bath for 30 min, before transferring them to a fresh plate seeded with 0P50 and incubated at 22.5°C

### Evaluation of the development stage of the animals

Developmental stage of wild type worms is evaluated 2 days post induction. Supposedly arrested animals were moved to new plates to assess the reality of the developmental arrest. For *mes-2* strains, F2 animals are sterile and develop slightly slower than wild-type, which allowed staging at day 3 post induction when they usually reach adulthood at 22.5°C. Worms were scored according to their size and the appearance of the vulva and gonad into L1, L2-3, L4 and adult. The *mes-2* worms were checked again at day 7 post induction to verify the results.

### Imaging and image analysis

The worms were imaged on an iMIC (FEI Munich GmbH) using the 10X air, 20X air, 40X oil, 60X 1.4 NA oil lens equipped with filters for DIC, brightfield *mCherry* and *GFP* detection and an ORCA-R2 CCD camera (Hamamatsu). To count cells expressing muscle-specific GFP::H2B, whole worms were imaged with 40x magnification in z stacks with a 1.5 μm distance between planes. Pictures were stitched together using Fiji and the number of GFP positive muscle cells was counted using point picker. To measure global *mCherry* expression, regions of interests were defined around animals and in the background to measure average fluorescence by animal. Final graphical assembly and comparisons were made in Microsoft Excel and R. All imaging experiments were performed at least twice with a minimum of 3 samples.

### Pharyngeal Pumping Measurements

Measurements of the pharyngeal pumping rate was performed as described^31^. Countings were performed for each worm 10 times at 30 seconds intervals in feeding conditions. If a worm crawled out of the food, counting was stopped until the worm moved again into OP50.

### RNAi experiments by feeding

Double stranded expressing bacteria Ahringer library, ^32^ were seeded on NG2 plates to deplete expression of the targeted genes. Heterozygous L3-L4 *mes-2* animals were moved onto RNAi plates and *mes-2* homozygous F2 generation was used for the experiments. After induction of the transcription factor expression, worms were moved to RNAi plates again.

Strain Genotype

**Strains list**
| Strain | Genotype |
| --- | --- |
| PMW4 | *gwls4[baf-1::GFP-lacI;myo-3::RFP] X* |
| PMW200 | *rrrSi261[myo-3p::cey-4::SL2::gfp:h2b; unc-119] I. ubSi13[hsp-16.2p::hlh-1::SL2::mCherry] II. unc-119(ed3) III.* |
| PMW217 | *rrrSi261 I. ubSi14[hsp-16.2p::end-1::SL2::mCherry] II. unc-119(ed3) III.* |
| PMW218 | *rrrSi261 I. gwSi3[hsp16.3p::mCherry] II. unc-119(ed3) III.* |
| PMW262 | *rrrSi261 I. ubSi13 II. unc-119(ed3) met-2(n4256) set-25(n5021) III.* |
| PMW283 | *rrrSi261 I. gwSi3 II. unc-119(ed3) met-2(n4256) set-25(n5021) III.* |
| PMW284 | *rrrSi261 I. mes-2(bn11) unc-4(e120) / mnC1 II.* |
| PMW285 | *rrrSi261 I. ubSi13 mes-2(bn11) unc-4(e120) / mnC1 II. unc-119(ed3) III.* |
| PMW292 | *ubSi13 II. unc-119(ed3) III. gwls4 X.* |
| PMW392 | *rrrSi261 I. ubSi14 mes-2(bn11) unc-4(e120) / mnC1 II. unc-119(ed3) III.* |
| PMW426 | *rrrSi261 I. mes-2(bn11) unc-4(e120) / mnC1 nls190 II.* |
| PMW427 | *rrrSi261 I. ubSi13 mes-2(bn11) unc-4(e120) / mnC1 nls190 II. unc-119(ed3) III.* |
| PMW428 | *rrrSi261 I. ubSi14 mes-2(bn11) unc-4(e120) / mnC1 nls190 II. unc-119(ed3) III.* |
| PMW476 | *rrrSi261 I. ubSi13 II. lin-12(n905) III.* |
| N2 | Wild-type Bristol strain |

## Supplementary Figures

**Figure 1-S1.**
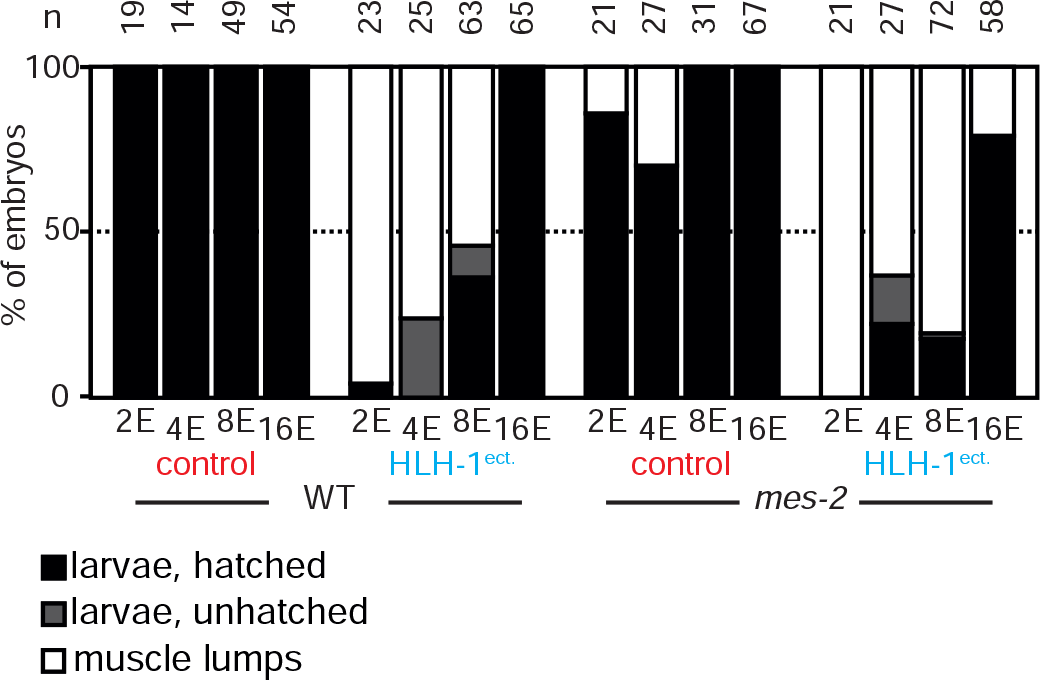
Phenotypic consequences of the ectopic induction of *hlh-1* or *mCherry* in wild-type and *mes-2* embryos from 2E, 4E, 8E and 16E stage, scored 24h post induction. The black bars indicate larvae that develop and hatch, the grey bar animals which developed but failed to hatch while the white bars are animals converted to muscle lumps.

**Figure 1-S2.**
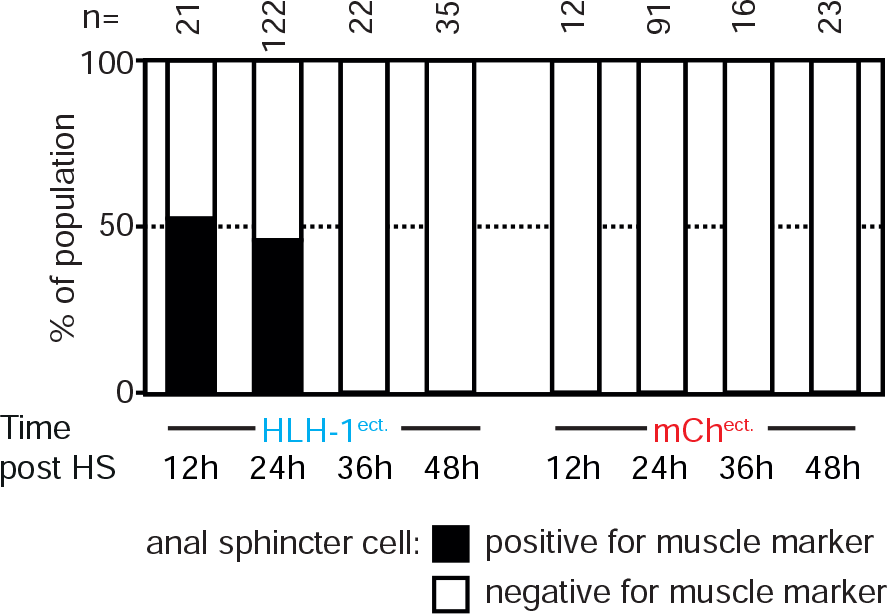
Proportion of the population in which the anal sphincter cell expresses the muscle marker in wild-type animals after ectopic induction of HLH-1 or *mCherry* expression at the L1 stage, scored 12, 24, 36 and 48 hours post-induction. The black bars represent the presence of *myo-3p*::H2B::GFP expression at the sphincter cell at a similar or stronger expression level than the weakest *bona fide* muscle cell in the tail region and the white bars represent an expression signal weaker than the surrounding muscle cells.

**Figure 1-S3.**
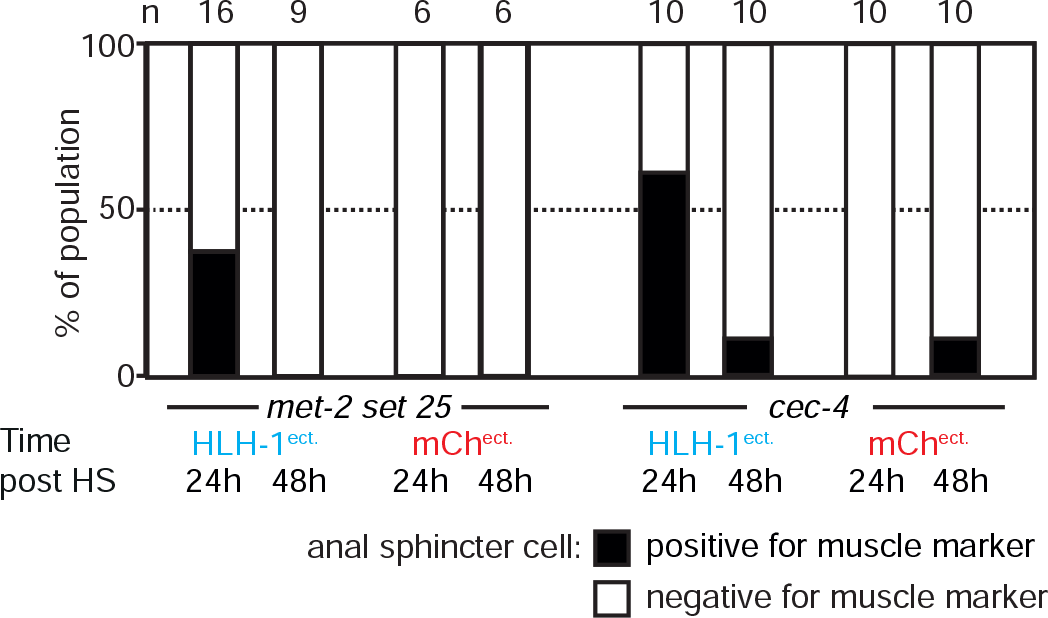
Proportion of the population in which the anal sphincter cell expresses the muscle marker in *set-25 met-2* or *cec-4* mutant animals after ectopic induction of HLH-1 or *mCherry* expression at the L1 stage, scored 24 and 48 hours post-induction. Scoring as in Figure 1-S2.

**Figure 3-S1.**
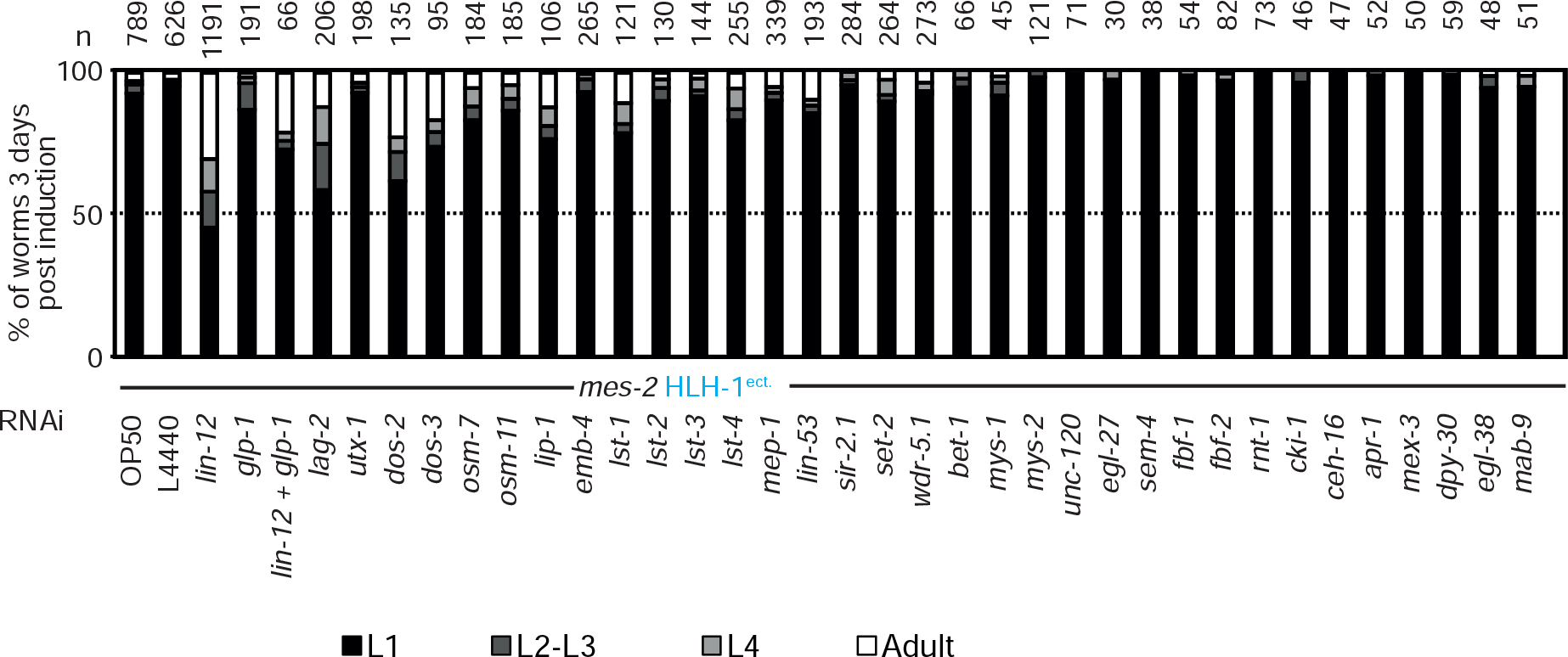
Evaluation of worm development stages 3 days post L1 HS of *mes-2* HLH-1 worms hatched after feeding parents on different RNAi conditions.

Table S1

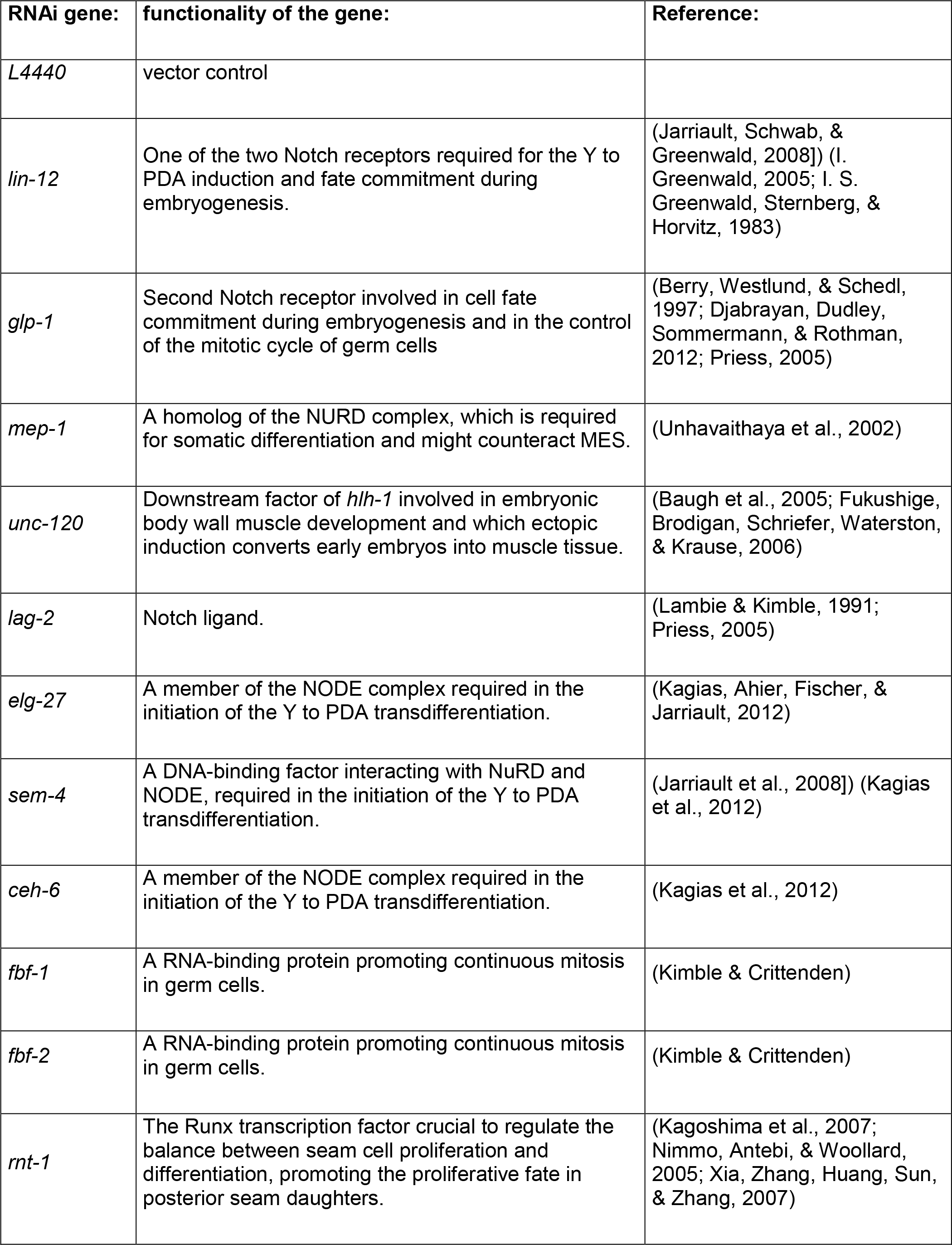

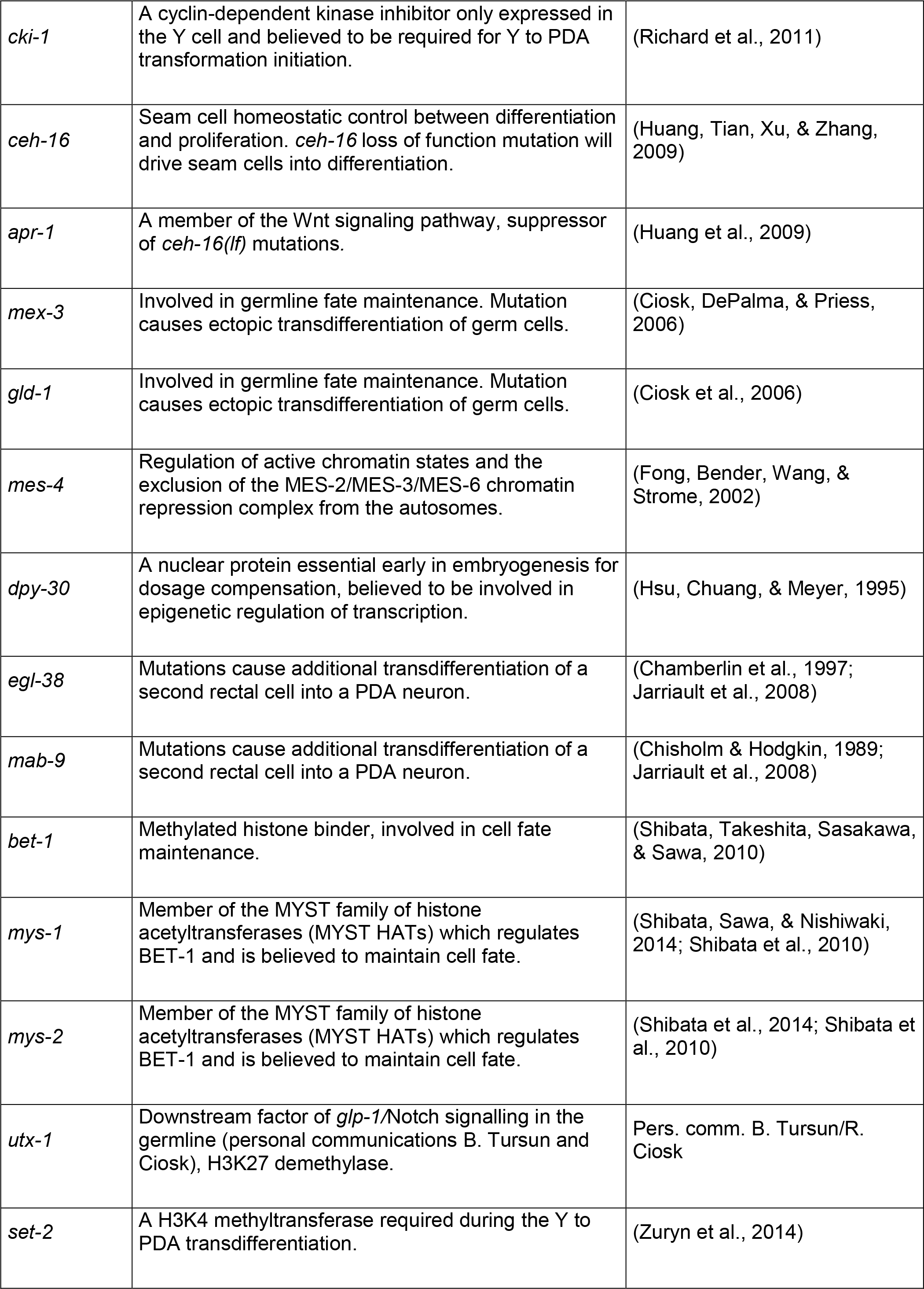

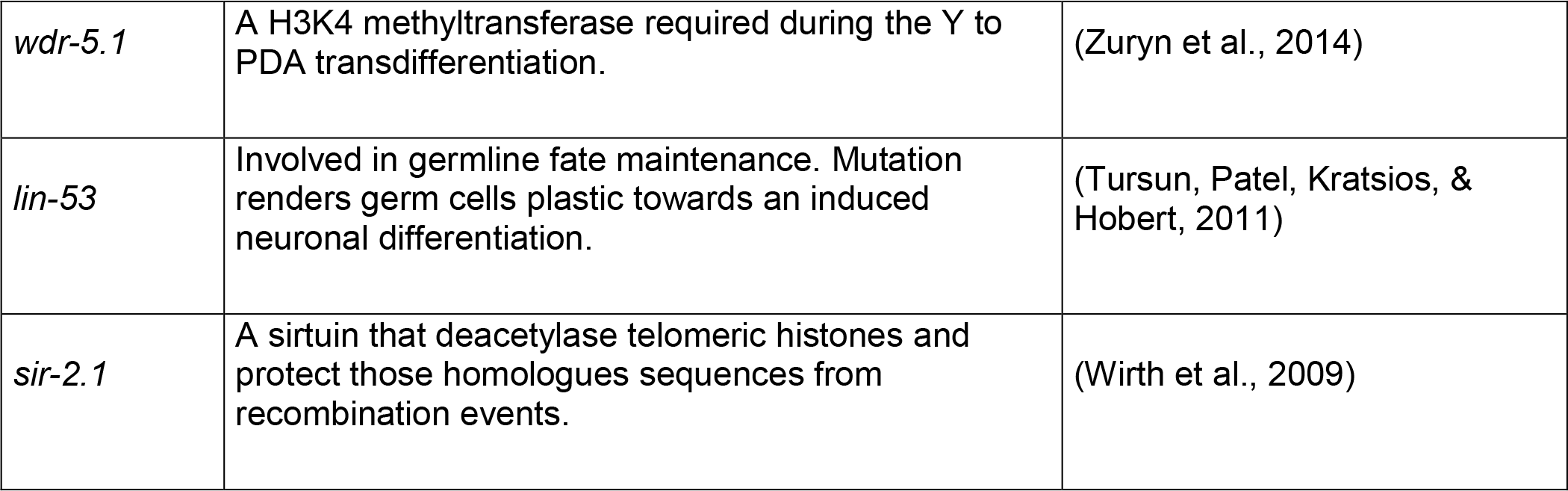

## Acknowledgements

The authors wish to thank members of the Meister laboratory for numerous discussions; Julie Campos for expert technical help. We wish to thank Dr. Rafal Ciosk for the muscle marker and Dr. Iskra Katic for help with CRISPR-mediated mutagenesis. Some strains were provided by the CGC, which is funded by NIH Office of Research Infrastructure Programs (P40 0D010440). The Meister laboratory is supported by the Swiss National Science Foundation (SNF assistant professor grant PP00P3_133744/159320), the Swiss Foundation for Muscle Diseases Research and the University of Bern.

